# SARS-CoV-2 spike protein interacts with and activates TLR4

**DOI:** 10.1101/2020.12.18.423427

**Authors:** Yingchi Zhao, Ming Kuang, Ling Zhu, Junhong Li, Zijing Jia, Xuefei Guo, Xiangxi Wang, Fuping You

## Abstract

The onset of sepsis is an important feature of COVID19 and a main cause of death. It is unknown how SARS-CoV-2 infection results in viral sepsis in human. We recently found that SARS-CoV-2 provoked an anti-bacterial like response and activation of TLR4 pathway at the very early stage of infection in animal models. This abnormal immune response led to emergency granulopoiesis and sepsis. However, the original trigger of TLR4 signaling by SARS-CoV-2 is unknown. We here identified that the trimeric spike protein of SARS-CoV-2 could bind to TLR4 directly and robustly activate downstream signaling in monocytes and neutrophils. Moreover, specific TLR4 or NFKB inhibitor, or knockout of MyD88 could significantly block IL-1B induction by spike protein. We thus reveal that spike protein of SARS-CoV-2 functions as a potent stimulus causing TLR4 activation and sepsis related abnormal responses.

## Introduction

Accumulating data obtained from numerous cohorts suggested that the main causes of death by COVID-19 include respiratory failure and the onset of sepsis (López-Collazo et al., 2020). Importantly, sepsis has been observed in nearly all deceased patients (Chao et al., 2020; Chen et al., 2020; Eastin and Eastin, 2020; Zhou et al., 2020). It is believed that the sepsis features are often related to the bacterial infection (Li et al., 2020). Moreover, age, procalcitonin and interleukin (IL)-6 levels, leukocytosis and lymphocytopenia have been included as factors associated with mortality in patients with COVID-19 (Zhou et al., 2020). Abnormalities of these factors in patients with unfavorable progression, combined with the high incidence of sepsis, strongly suggest the dysregulation of the host’s immune response. Toll like receptor 4 (TLR4) is the pattern recognition receptor that mediates anti-gram negative bacterial immune responses by recognizing Lipopolysaccharide (LPS) from bacteria (Poltorak et al., 1999). We recently found that SARS-CoV-2 infection provoked an anti-bacterial like response at the very early stage of infection in rhesus macaque and hACE2-transgenic mice models. The S100A8/A9-TLR4 are the critical host factors that result in emergency granulopoiesis and sepsis in the SARS-CoV-2 infected animals. However, it is unknown what is the original trigger initiating these abnormal immune responses during SARS-CoV-2 infection.

We here identified that the spike protein of SARS-CoV-2 was able to interact with TLR4 directly and activated the production of proinflammatory cytokines. MyD88 and NFKB were required for IL-1B induction by spike protein. The spike protein thus is the original stimulus provoking activation of TLR4 signaling pathway.

## Results

### Spike protein of SARS-CoV-2 interacted with TLR4 directly

Previous silico studies predicted that cell surface TLRs, especially TLR4 are most likely to be involved in recognizing molecular patterns, probably spike protein, from SARS-CoV-2 to induce inflammatory responses (Bhattacharya et al., 2020; Choudhury and Mukherjee, 2020). Combined with our recent data that TLR4 signaling was activated by SARS-CoV-2 infection, we hypothesized that spike protein of SARS-CoV-2 could activated TLR4 pathway. A recent study showed that trimeric SARS-CoV-2 spike proteins are high quality antigens (Pino et al., 2020). To this end, we purified the trimeric spike protein (1-1208aa) (Figure 1A and 1B), as this form of spike protein presents on the surface of viral particle, which most likely interacted with the proteins on the surface of host cells. The surface plasmon resonance (SPR) assay showed that SARS-CoV-2 spike trimer directly bound to TLR4 with a high affinity (Figure 1C).

**Figure 1.**
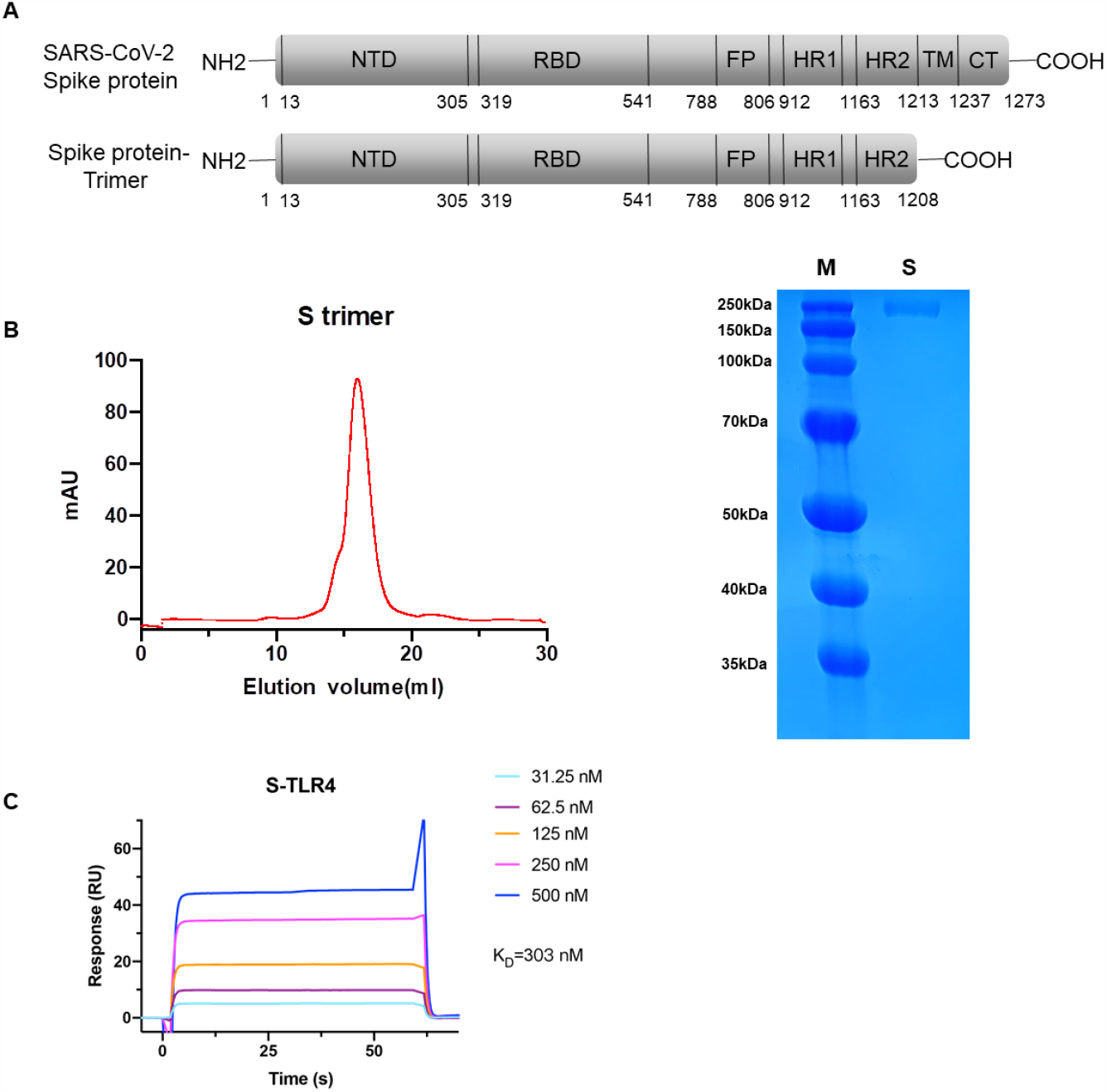
Spike protein of SARS-CoV-2 interacted with TLR4 directly. (A) Schematic representation of full-length-spike protein and trimer-ectodomain-spike protein of SARS-CoV-2. (B.C) Affinity analysis of the binding of TLR4 to SARS-CoV-2 S trimer. Purified SARS-CoV-2 S trimer was immobilized onto a CM5 sensor chip surface and tested for real-time association and dissociation of the TLR4.

### Spike protein induces the production of proinflammatory cytokines

To evaluate its function, we treated THP-1 cell, human monocyte derived from patient with acute monocytic leukemia, with purified spike protein. qRT-PCR analysis showed that IL-1B was robustly induced by spike protein, which was comparable to the positive control LPS (Figure 2A). To confirm this, we treated cells with an increasing amount of spike protein and found that IL-1B expression was increased in a dose dependent manner (Figure 2B and 2C). Moreover, IL-6 was also induced by spike protein. As IL-1B induction was much more significant than that of IL-6 (Figure 2D), we then chose IL-1B production as a marker of immune activation. More interestingly, the pseudo virus with spike protein was also able to induce IL-1B production (Figure 2E). In addition to monocyte, neutrophils are important myeloid cells in sepsis with high expression of TLR4 on their cell surface. We utilized all-trans retinoic acid (ATRA) to treat HL-60 cell, a promyelocytic leukemia cell line, which directed those cells to differentiate into neutrophils. Spike proteins significantly induced IL-1B production in HL-60 cells after ATRA treatment (Figure 2F and 2G).

**Figure 2.**
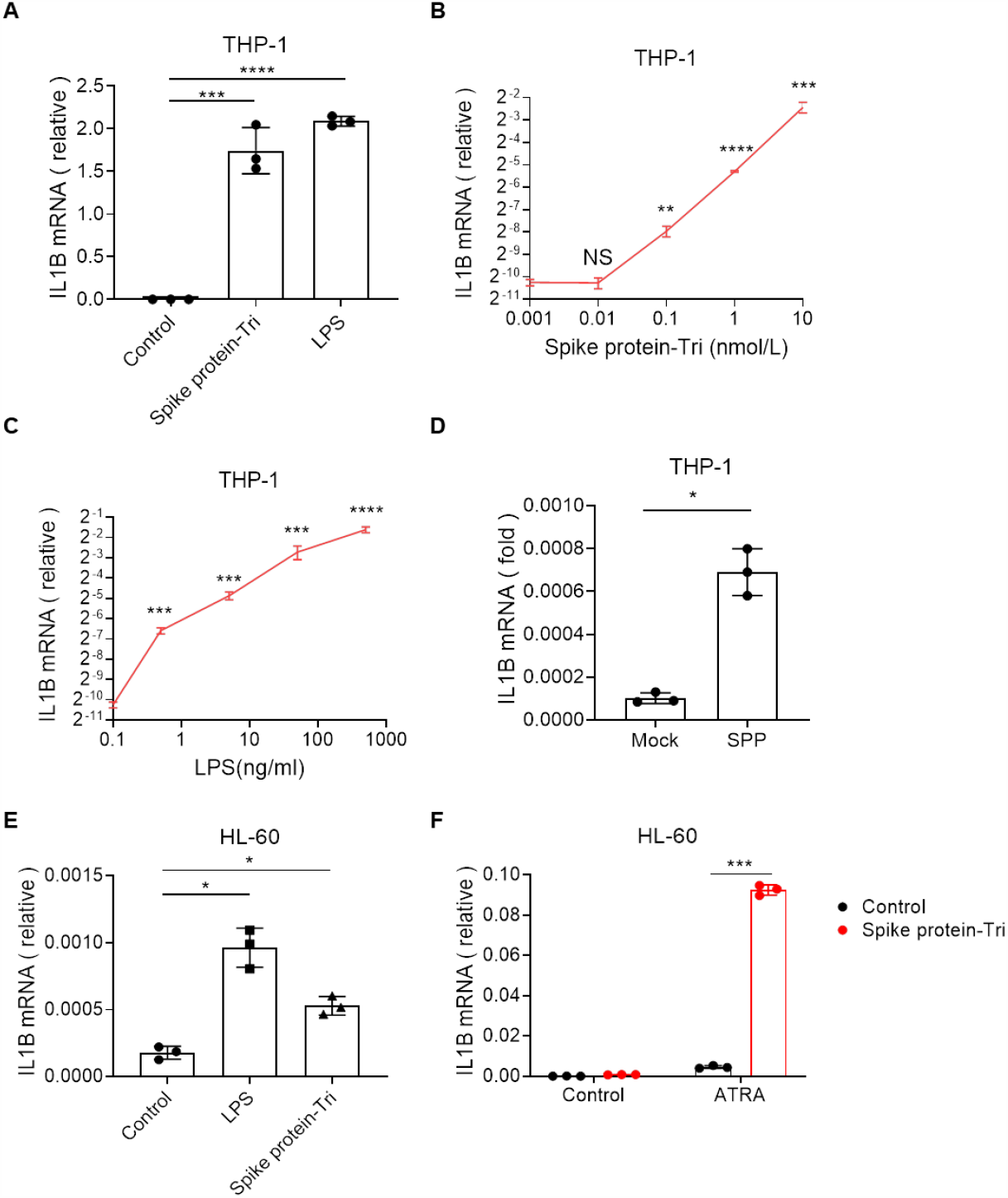
Spike protein induces the production of proinflammatory cytokines. (A) qRT-PCR analysis for the expression of IL1B in the THP-1 cells treated with control, 500 ng/ml LPS, and 10 nM Spike protein-EC for 12 hours. n = 3. (B) qRT-PCR analysis for the expression of IL1B in the THP-1 cells treated with control, 0.01 nM, 0.1 nM, 1 nM, 10 nM Spike protein-EC for 12 hours. n = 3. (C) qRT-PCR analysis for the expression of IL1B in the THP-1 cells treated with control, 0.5 ng/ml, 5 ng/ml, 50 ng/ml, 500 ng/ml LPS for 12 hours. n = 3. (D) qRT-PCR analysis for the expression of IL1B in the THP-1 cells treated with control and Spike protein-pseudotyped (SPP) lentivirus for 12 hours. n = 3. (E) qRT-PCR analysis for the expression of IL1B in the HL-60 cells treated with control, 500 ng/ml LPS, and 10 nM Spike protein-EC for 12 hours. n = 3. (F) qRT-PCR analysis for the expression of IL1B in the Control group and ATRA-differentiated HL-60 cells treated with 1 nM Spike protein-EC for 12 hours. n = 3. (NS=non-significant, *P < 0.05; **P < 0.01; ***P < 0.001; ****P < 0.0001)

### Spike protein activates inflammation via TLR4 pathway

To gain further insight of the mechanism by with spike protein activates TLR4 signaling, we treated THP-1 cells with the N-terminal domain (NTD) and the receptor-binding domain (RBD) truncated spike protein. Only the trimer form protein can induce IL-1B and IL-6 (Figure 3A and 3B). To examine if this activation was mediated by TLR4, we treated cells with TLR4 inhibitor, Resatorvid. Resatorvid treatment greatly blocked induction of IL-1B by spike protein and LPS. Moreover, spike protein was also able to induce IL-1B production in murine macrophage cell line (Raw 264.7) in a TLR4 and MyD88 dependent way (Figure 3E). The NF-KB (JSH-23) but not ACE2 (MLN-4760) inhibitor was able to suppress IL-1B induction by spike protein (Figure 3F and 3G) suggesting that ACE2 was not involved in TLR4 signaling activation by spike protein. Trypsin digestion almost completely abolished the activation of IL-1B by spike protein ruling out the possibility that the protein was contaminated by LPS.

**Figure 3.**
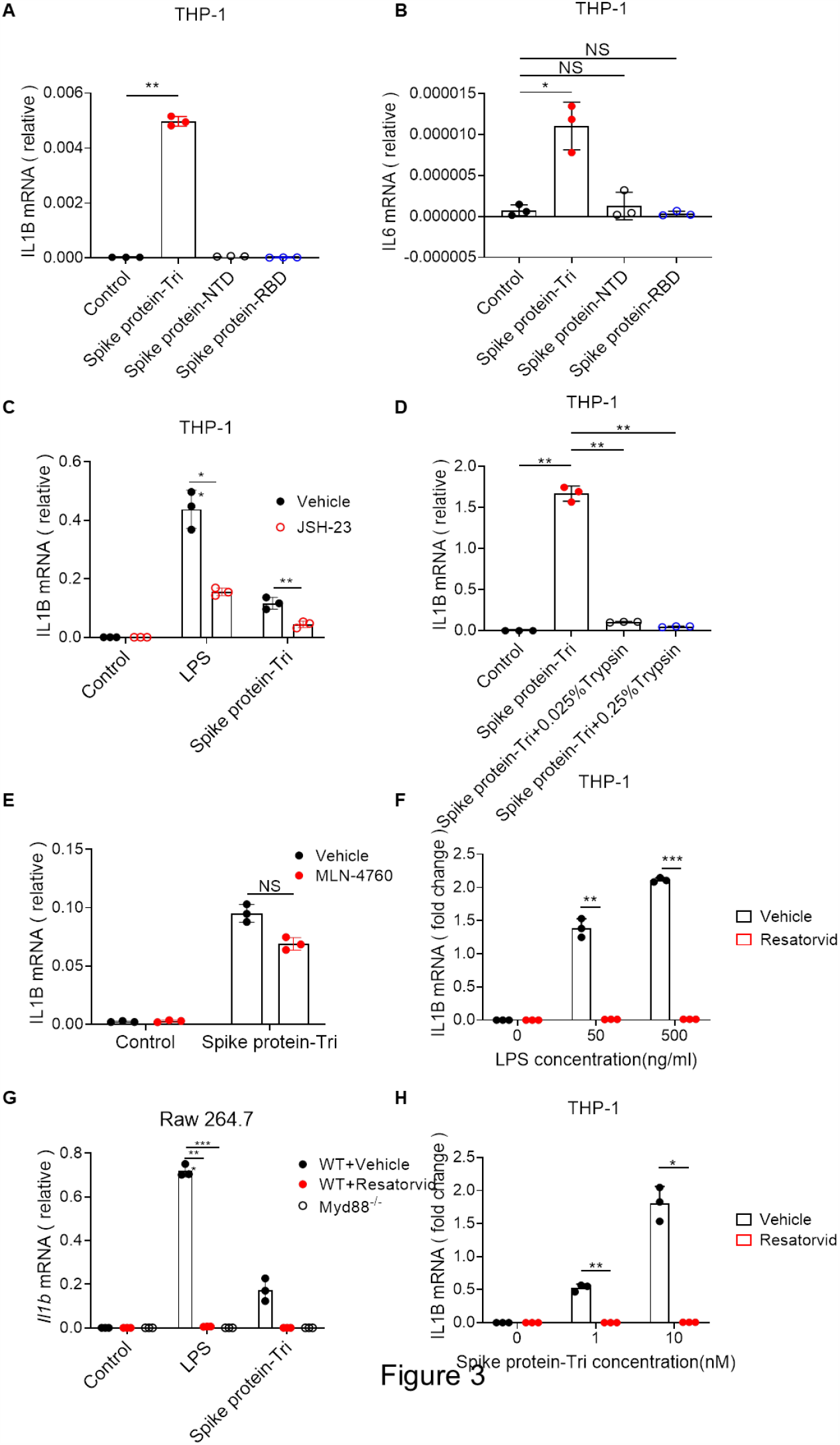
Spike protein activates inflammation via TLR4 pathway. (A) qRT-PCR analysis for the expression of IL1B in the THP-1 cells treated with Ectodomain (EC), N-terminal domain (NTD), Receptor binding domain (RBD) of SARS-CoV-2 Spike protein at 10 nM, and control for 12 hours. n = 3. (B) qRT-PCR analysis for the expression of IL6 in the THP-1 cells treated with Ectodomain (EC), N-terminal domain (NTD), Receptor binding domain (RBD) of SARS-CoV-2 Spike protein at 10 nM, and control for 12 hours. n = 3. (C) qRT-PCR analysis for the expression of IL1B in the THP-1 cells treated with control, 500 ng/ml LPS, and 10 nM Spike protein-EC for 12 hours with or without 50 uM JSH-23 treatment. n = 3. (D) qRT-PCR analysis for the expression of IL1B in the THP-1 cells treated with control, 10 nM Spike protein-EC, 10 nM Spike protein-EC + 0.025% Trypsin, 10 nM Spike protein-EC + 0.025% Trypsin for 12 hours. n = 3. (E) qRT-PCR analysis for the expression of IL1B in the THP-1 cells treated with control, 1 nM Spike protein-EC for 12 hours with or without 10 μM MLN-4760 treatment. n = 3. (F) qRT-PCR analysis for the expression of IL1B in the THP-1 cells treated with control, 50 ng/ml LPS, and 500 ng/ml LPS for 12 hours with or without 100 μM Resatorivd treatment. n = 3. (G) qRT-PCR analysis for the expression of *Il1b* in the WT and Myd88^-/-^ Raw 264.7 cells treated with control, 500 ng/ml LPS, 10 nM Spike protein-EC for 12 hours with or without 1 μM Resatorvid treatment. n = 3. (H) qRT-PCR analysis for the expression of IL1B in the THP-1 cells treated with control, 1 nM Spike protein-EC, and 10 nM Spike protein-EC for 12 hours with or without 100 μM Resatorivd treatment. n = 3. (NS=non-significant, *P < 0.05; **P < 0.01; ***P < 0.001)

### MHV-A59 activates TLR4 pathway

To determine if other coronavirus could activate TLR4 signaling, we treated THP-1 cells with murine coronavirus MHV-A59. Interestingly, MHV-A59 significantly induced IL-1B in a dose dependent manner (Figure 4A). The induction could be blocked by TLR4 inhibitor (Figure 4B) indicating that MHV-A59 activated TLR4 pathway. Theoretically, there is no MHV-A59 receptor (murine *Ceacam1*) expression in THP-1 cells, so MHV-A59 was not able to infect and enter this human monocyte. To confirm this, we washed those cells after treatment. After washing with PBS, the viral load was significantly decreased, so was the induction of IL-1B (Figure 4). These data suggested that MHV-A59 could trigger TLR4 signaling probably via spike-TLR4 interaction.

**Figure 4.**
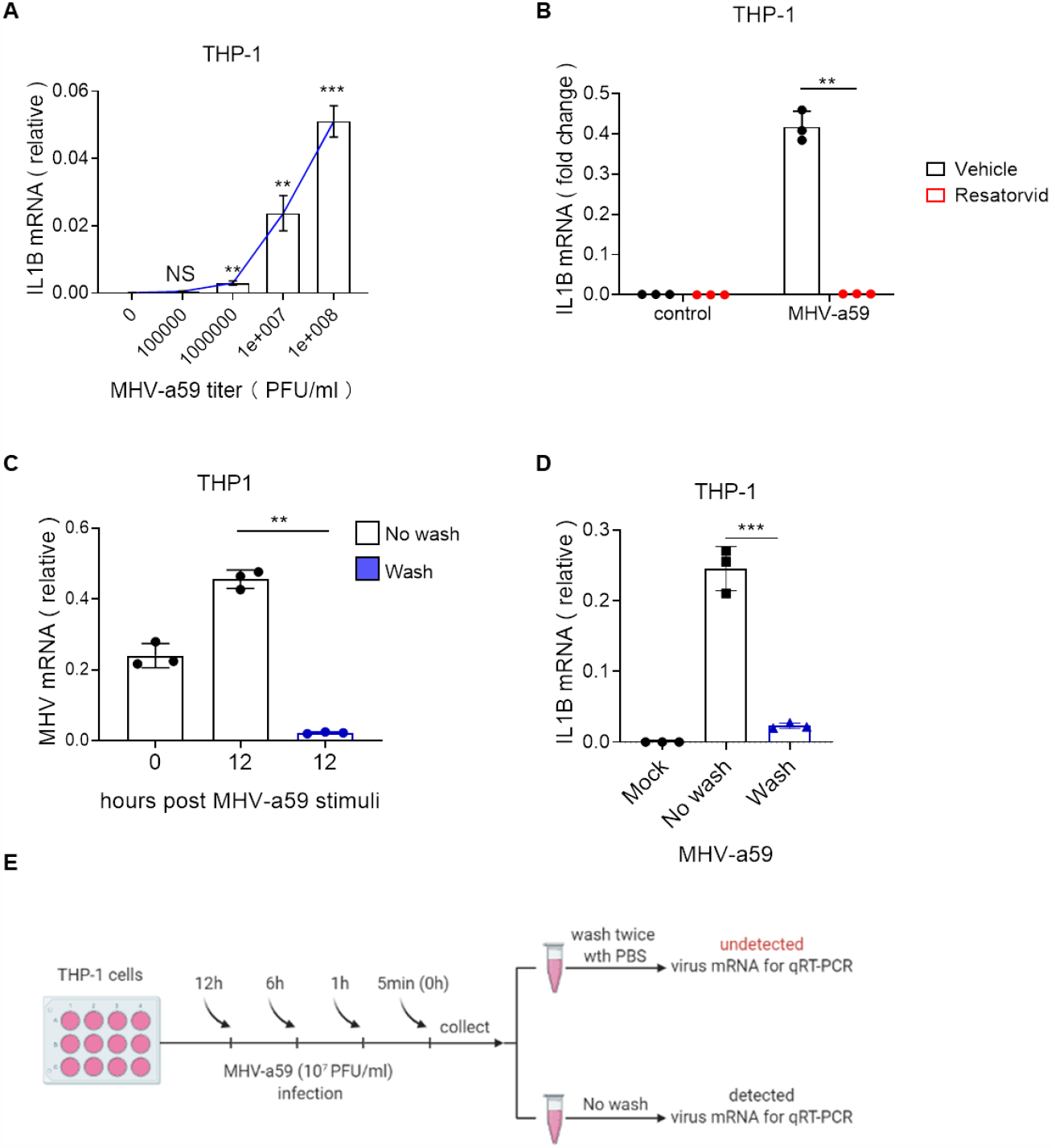
MHV-A59 activates TLR4 pathway. (A) qRT-PCR analysis for the expression of IL1B in the THP-1 cells treated with 1×10^7^ PFU/ml MHV-A59 for 0 and 12 hours with or without washing by PBS. n = 3. (B) qRT-PCR analysis for the expression of IL1B in the THP-1 cells treated with control, 1×10^5^ PFU/ml, 1×10^6^ PFU/ml, 1×10^7^ PFU/ml, 1×10^8^ PFU/ml MHV-A59 for 12 hours. n = 3. (C) qRT-PCR analysis for the MHV-A59 titer in the THP-1 cells treated with 1×10^7^ PFU/ml MHV-A59 for 0 and 12 hours with or without washing by PBS. n = 3. (D) qRT-PCR analysis for the expression of IL1B in the THP-1 cells treated with 1×10^7^ PFU/ml MHV-A59 for 12 hours with or without 100 μM Resatorivd treatment. n = 3. (E) A flow chart depicting the process of MHV-A59 stimulating THP-1 cells with or without washing.

## Discussion

The genome of SARS-CoV-2 is positive single strand RNA. A mount of double strands RNA (dsRNA) is generated during SARS-CoV-2 replication. dsRNA is an important pathogen associated molecular pattern (PAMP) that can be recognized by host pattern recognition receptors (PRRs), including MDA5, RIG-I and TLR3. Subsequently, those PRRs transduce danger signals to downstream pathways and trigger host defense against invading virus, particularly the production of anti-viral effectors, such as type I IFNs. However, type I IFNs induction is completely blocked during SARS-CoV-2 infection (Shuai et al., 2020). Proinflammatory cytokines, such as IL-1B and IL-6, were significantly induced by SARS-CoV-2 infection in patients. These data indicated the SARS-CoV-2 infection caused dysregulation of immune responses. To this end, we recently examined the immune responses at the early stage of SARS-CoV-2 infection in animal models (Guo et al., 2020). Instead of typical anti-viral immune responses, anti-bacterial responses were triggered by SARS-CoV-2 at day 1 after infection. Alarmin S100A8/9 and TLR4 axis promoted emergency granulopoiesis and expansion of premature neutrophils, which was a typical feature of systemic bacterial infection and sepsis. All these serinio were mirrored in clinical observations of COVID-19 patients. For example, the number of premature neutrophils was significantly increased in the COVID-19 patients with severe symptoms compared to healthy control or asymptomatic patients. Elevated S100A8/9 and neutrophils was positive relevant with thrombosis in COVID-19 patients. Together, we could reason that the abnormal early immune responses contributed to unfavorable clinical features of COVID-19 patients. We here found that, before entering the cells, the viral surface spike protein interacted with and activated TLR4. Thus, an anti-bacterial response was initiated at the very early stage of SARS-CoV-2 infection.

The SPR assay showed that trimeric spike protein interacted with TLR4 directly. However, it is unknown which domain or motif mediates their interaction. We evaluated the induction of IL-1B and IL-6 by trimeric spike protein, which was comparable to LPS treatment. It is noteworthy to investigate if spike protein triggers similar or identical immune responses to LPS treatment. Moreover, MHV-A59 activated TLR4 signaling without entering cell indicating that the spike protein of MHV-A59 was also able to activate TLR4. Therefore, future work should address if it is the common ability of spike proteins from different types of coronaviruses.

## Acknowledgments

This work was supported by the National Natural Science Foundation of China (31570891; 31872736), the National Key Research and Development Program of China (2016YFA0500300; 2020YFA0707800), the National Key Research and Development Program (2020YFA0707500) and the Strategic Priority Research Program (XDB29010000). Xiangxi Wang was supported by Ten Thousand Talent Program and the NSFS Innovative Research Group (81921005).

## Author Contributions

F.Y., X.W. and Y.Z. conceived the study and analyzed the data. Y.Z., M.K. L.Z. and J.L. performed most experiments and analyzed the data. Z.J..and X.<u>G.</u> provided support on literature search. F.Y. wrote the paper. F.Y. and X.W. revised the paper.

## STAR Methods

### RESOURCE AVAILABILITY

#### Lead Contact

Further information and requests for resources and reagents should be directed to and will be fulfilled by the Lead Contact, Fuping You.

#### Materials Availability

This study did not generate new unique reagents.

#### Data and Code Availability

This study did not generate high throughput data.

### EXPERIMENTAL MODEL AND SUBJECT DETAILS

#### Cell culture

293T cells, Raw 274.7 cells, 17CL-1 cells, THP-1 cells and HL-60 cells were kept in our lab. 17CL-1 cells were cultured in DMEM medium (Gibco) supplemented with 10% FBS (PAN), 100 U/mL Penicillin-Streptomycin. Raw 274.7 cells, THP-1 cells and HL-60 cells were cultured in 1640 medium (Gibco) supplemented with 10% FBS, 100 U/mL Penicillin-Streptomycin. Cells were negative for mycoplasma. Isolation of BMDMs (bone-marrow derived macrophages) and peritoneal macrophages was performed as described (Barber, 2009).

#### Viruses

MHV-A59 (mouse hepatitis virus A-59) has been described previously (Yang et al., 2014). MHV-A59 were propagated in 17CL-1 cells followed by 3 cycles of freezing and thawing. The large debrises were spun down and the supernatants were ultrafiltered and concentrated by 100 KD ultrafiltration device (Millipore). The supernatants of ultrafiltration device were used as a stock solution. The titer of the viruses was determined by plaque assay in 17CL-1 cells.

### METHOD DETAILS

#### Expression constructs

The plasmids used for protein expression were constructed by insertion of the coding sequences for SARS-CoV-2 S trimer (residues 1–1208, GenBank:MN908947.3) into the mammalian expression vector pCAGGS with a C-terminal twin Strep tag to facilitate protein purification. The S protein gene was constructed with proline substitutions at residues 986 and 987, a “GSAS” instead of “RRAR”at the furin cleavage site according to Jason S. McLellan”s research.

#### Protein expression and purification

Expi293F cells (Thermo Fisher) were transiently transfected with the S protein expression construct using polyethylenimine. To purify the S trimer protein, filtered cell supernatants was loaded on a Strep-tactin resin (IBA). The column was then washed with 5 column volumes of Buffer W (100 mM Tris-HCl, pH 8.0, 150 mM NaCl, 1 mM EDTA). The S trimer was eluted by Buffer W containing 50 mM biotin. Elution fractions were analyzed by SDS-PAGE. The protein was subjected to additional purification by gel filtration chromatography using a Superose 6 10/300 column (GE Healthcare) in 20 mM Tris, 200 mM NaCl, pH 8.0.

#### Surface plasmon resonance (SPR)

For the binding affinity assay, purified SARS-CoV-2 S trimer was immobilized onto a CM5 sensor chip surface by using the NHS/EDC method to a level of ∼ 700 response units (RUs) using Bia-core 8000 (GE Healthcare) and a PBS running buffer (supplemented with 0.05% Tween-20) was prepared for assay. Serial dilutions of purified TLR4 were injected. The sample flew over the chip at a rate of 20 μl/min for 30 s, then the dissociation of the sample was at the same rate for another 30 s. All antibodies were regenerated with Gly 1.5. The response of the sample to the S trimer was recorded at room temperature and the data was analyzed by Bia-core 8000 Evaluation Software (GE Healthcare).

#### Generation of pseudotyped lentivirus

SARS-CoV-2 S gene (GenBank: QHU36824.1) fusion with a C-terminal 3xFlag tag was synthesized and cloned into a pMD2.G vector. 293T cells were grown in DMEM containing 10% FBS and co-transfected by pCDH-eGFP (6000 ng), psPAX2 (2000 ng) and pMD2.G-Spike (2000 ng) or pMD2.G-VSVG (2000 ng). The supernatant with produced virus (Spp or VSV-G lentivirus) was harvested 48-hours post transfection, clarified by centrifuging at 8000 g for 10 min at 4^°^C. The virus was collected by an ultracentrifugation at 50000 g for 2 hours (hrs) using Beckman SW41 rotor. The viral pellets were resuspended by FBS free 1640 medium and stored at -80°C before use. The viral particle number was determined using a real time RT-PCR assay to quantify the RNA copies of eGFP.

#### HL-60 cells differentiation

HL-60 cells were cultured in 1640 medium (Gibco) with 10% FBS in 5% CO2 humidified air at 37°C. Cells were passaged every 3 days and only cells passaged no more than 15 times were used for all experiments. Differentiation of HL-60 cells into granulocyte-like cells was performed as described (Manda-Handzlik et al., 2018). The cells were incubated for 5 days with ATRA (1 μ mol L^-1^). After 5 days of differentiation, cells were collected into a 15 ml tube and precipitated naturally for 2 hours. Cell pellet was resuspended by 1640 medium with 10% FBS. Differentiated cells were counted before experiment.

#### Quantitative RT-PCR (qRT-PCR) analysis

Total RNA was isolated from the tissues by TRNzol reagent (DP424, Beijing TIANGEN Biotech, China). Then, cDNA was prepared using HiScript III 1st Strand cDNA Synthesis Kit (R312-02, Nanjing Vazyme Biotech, China). qRT-PCR was performed using the Applied Biosystems 7500 Real-Time PCR Systems (Thermo Fisher Scientific, USA) with SYBR qPCR Master mix (Q331-02, Nanjing Vazyme Biotech, China). The data of qRT-PCR were analyzed by the Livak method (2^−ΔΔCt^). Ribosomal protein L19 (RPL19) was used as a reference gene for mouse cell line, and GAPDH for human cell line. qRT-PCR primers are displayed in supplementary materials Table S1.

### QUANTIFICATION AND STATISTICAL ANALYSIS

#### Statistical analysis

All analyses were repeated at least three times, and a representative experimental result was presented. Two-tailed unpaired Student’s t test was used for statistical analysis to determine significant differences when a pair of conditions was compared. Asterisks denote statistical significance (*P < 0.05; **P < 0.01; *** P < 0.001). The data are reported as the mean ± S.D.

